# Ophthalmologic findings in an induced model of Holoprosencephaly in zebrafish

**DOI:** 10.1101/2024.08.09.607329

**Authors:** Johannes Bulk, Valentyn Kyrychenko, Stephan Heermann

## Abstract

Early forebrain development is a fascinating process. The fate of brain function but also the fate of visual perception largely depends on it. Holoprosencephaly (HPE) is the most frequent developmental disorder of the forebrain, during which the separation of the early precursor domains is hampered. A spectrum of clinical manifestations is seen with severe forms like alobar HPE and less severe forms like lobar HPE. The ophthalmologic findings which accompany HPE are also found as a spectrum that ranges from ocular hypotelorism and synophthalmia to cyclopia and anophthalmia. Here we ask, whether anophthalmia or cyclopia is the default ophthalmologic finding in severe forms of HPE. In this brief analysis, we made use of a recently established zebrafish model of severe HPE, based on BMP ligand induction. Such BMP ligand induction resulted in anophthalmia. We attenuated the induction protocol to investigate whether the anophthalmia phenotype could be changed into a cyclopic phenotype. We found a spectrum of ocular phenotypes, ocular hypotelorism, and also cases of synophthalmia and cyclopia. This suggests that in the context of this HPE model the strongest ophthalmologic phenotype is anophthalmia and less severe forms are cyclopia, synophthalmia and ocular hypotelorism.

## Introduction

Holoprosencephaly (HPE) is the most frequent developmental pathology of the forebrain (Malta et al., 2023; Matsunaga and Shiota, 1977; Pineda-Alvarez et al., 2011) in which the splitting of precursor domains of the Anterior Neural Plate (ANP) is hampered. It is mostly caused by genetic defects, however, also environmental risk factors/ triggers are described (Addissie et al., 2021; Roessler et al., 2018).

Not all patients who suffer from HPE show the same specific phenotype but rather a spectrum of phenotypes. This spectrum was divided into groups of severe forms e.g. alobar HPE, intermediate forms, semilobar HPE, and less severe forms e.g. lobar HPE (Gomez et al., 2024). Both, the presumptive telencephalic lobes as well as both retinae derive from the ANP. Thus, it is consistent that ophthalmologic phenotypes are associated with HPE (Pineda-Alvarez et al., 2011). Such ophthalmologic phenotypes are also found as a spectrum. This ranges from coloboma, ocular hypotelorism, synophthalmia to cyclopia and anophthalmia (Fallet-Bianco, 2018; Pineda-Alvarez et al., 2011). Fundamental discoveries regarding the role of BMP signaling during early forebrain and eye development have been made in the mouse and chicken model. It was found in a mouse double Knock out of two BMP antagonists, Noggin and Chordin, that the formation of the forebrain was affected variably which resulted in e.g. cyclopia and aprosencephaly (Bachiller et al., 2000; Klingensmith et al., 2010). A Knock out of Noggin alone resulted in a milder form of HPE (Lana-Elola et al., 2011) and a double Knock out of two BMP receptors, Bmpr1a and Bmpr1b, affected dorsal aspects of the developing midline accompanied by microphthalmia and a new class of HPE was proposed (Fernandes et al., 2007). In chicken, BMP application via soaked beads resulted in cyclopia (Golden et al., 1999). We used zebrafish as a model and recently found that BMP antagonism is crucial for various consecutive steps of eye development. *Nog2, chrd, fsta* and *grem2b* were found expressed in or adjacent to the ANP and an induction of the BMP ligand *bmp4* at 8.5 hpf was sufficient to hamper ANP splitting which resulted in a severe form of HPE and anophthalmia (Bulk et al., 2023). In consecutive stages of development, we found *fsta* expressed in the optic vesicle and optic cup and inductions of *bmp4* at different stages of development hampered optic fissure formation and optic vesicle to optic cup transformation resulting in “morphogenetic coloboma” (Eckert et al., 2019; Heermann et al., 2015). At an even later stage of development, we found *grem2b* and *fsta* expressed in in the margins of the optic fissure. An induction of *bmp4* at this stage, resulted in a “fusion-inhibited coloboma” (Knickmeyer et al., 2018). Based on our findings, we propose a continuous gradient of these forebrain and eye developmental processes, all being crucially facilitated or protected by BMP antagonism. Disturbances of this BMP antagonism could explain well different ophthalmologic phenotypes associated with HPE including also coloboma. In this brief study we ask, whether anophthalmia or cyclopia is the default ophthalmologic phenotype in severe forms of HPE. We made use of the beforementioned model of a *bmp4* induction in zebrafish where an induction at 8.5 hpf resulted in anophthalmia (Bulk et al., 2023). Importantly, retinal precursors were found stuck in the forebrain as a “crypt-oculoid” (Bulk et al., 2023). Now, we attenuated the induction protocol. Our aim was to investigate whether the anophthalmia phenotype could be changed into less severe ophthalmologic phenotypes. We found ocular hypotelorism, synophthalmia and cases of cyclopia.

## Results

### Attenuated *bmp4* induction leads to graded eye phenotypes including cyclopia

We use a transgenic zebrafish line which expresses *bmp4* after heat-shock induction (*Tg(hsp70l:bmp4*,*myl7:EGFP*). This line was established and used previously (Bulk et al., 2023; Knickmeyer et al., 2018). Heat-shock induction at 8.5 hpf for 15 minutes resulted in a severe form of HPE (Bulk et al., 2023). The HPE phenotype was accompanied by anophthalmia regularly. Notably, retinal precursor cells were found inside the forebrain, a “crypt-oculoid”.

We now attenuated the *bmp4* induction to address whether the phenotype “anophthalmia” could be changed into e.g. less severe ocular phenotypes. We performed heat shocks at 8.5 hpf which were shorter in duration (**Figure 1, scheme of experimental design**) compared to the protocol which we used previously (Bulk et al., 2023). We induced *bmp4* expression for 10 minutes (100%, 186 embryos, with *cmlc2:GFP* expression were phenotypic, 100%, 198 embryos, which did not express cmlc2:GFP were not phenotypic) and 7.5 minutes (100%, 40 embryos with *cmlc2:GFP* expression were phenotypic, 97%, 36 embryos which did not express cmlc2:GFP were not phenotypic, 3%, 1 embryo which did not express cmlc2:GFP, showed ocular hypotelorism and coloboma). A summary of metadata can be found in the supplemental table. The induction for 10 minutes frequently resulted in a spectrum of phenotypes. Its range lead from synophthalmia, an incomplete separation of the eyes with yet two distinguishable optic vesicles (**Figure 1B-D, please see H-J as control**), to cyclopia (**Figure 1B’-D’, please see H-J as control**). We observed a continuous distribution of phenotypes in-between those two parameter values. Of a total of 186 fish observed, we can account for at least 7 synophthalmic and 3 entirely cyclopic fish. The latter displayed a severe pericardial effusion as in the HPE/anophthalmia phenotype previously described. The induction for 7.5 minutes resulted in less severe phenotypes, ocular hypotelorism, a reduced distance between the eyes (**Figure 1E-G, please see H-J as control)**.

**Figure 1.**
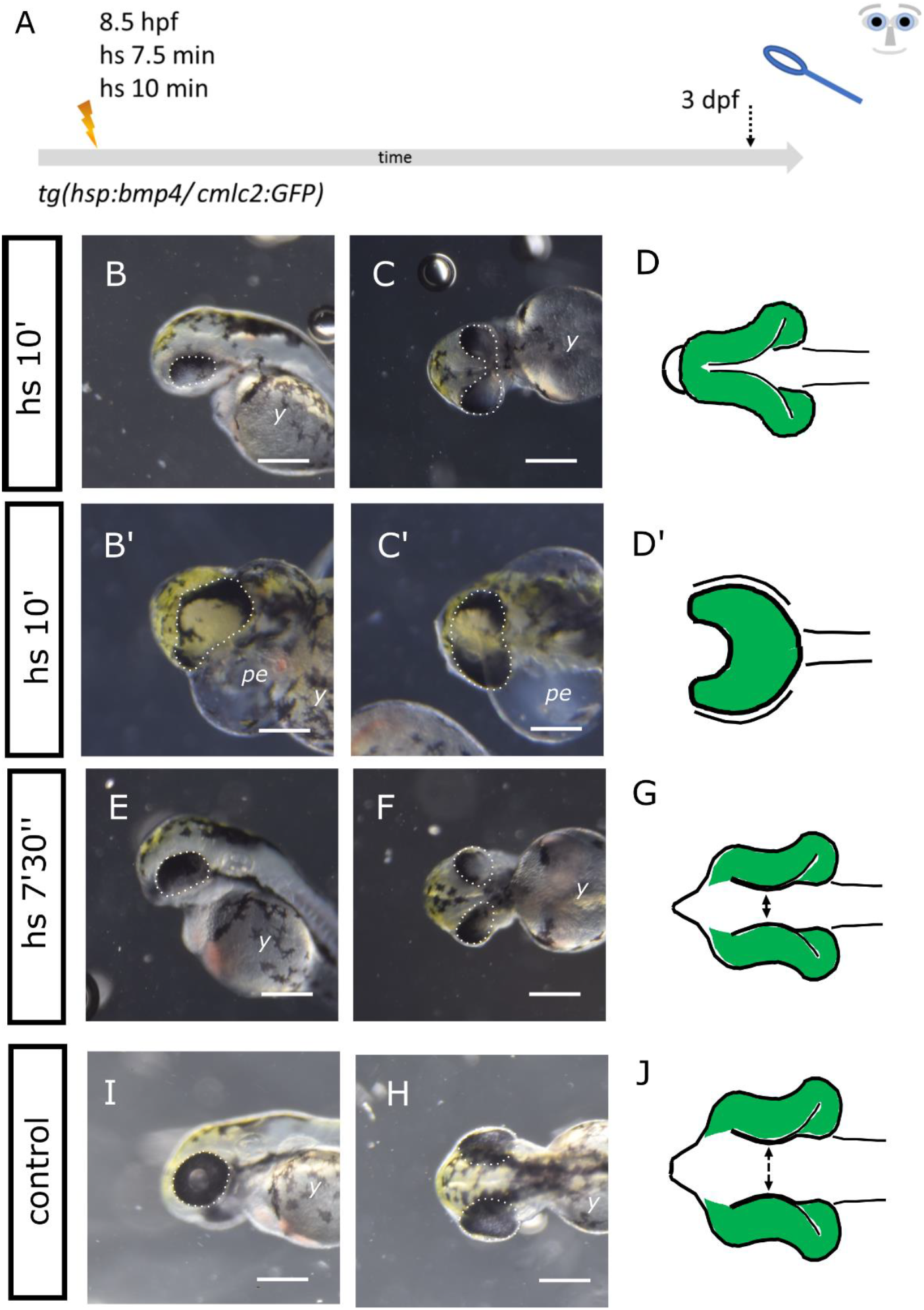
Attenuated *bmp4*-induction. A: summary of experimental procedure: embryos were heat-shocked at 8,5 hpf for 7,5 or 10 min and analyzed 3 dpf. B-D’: *bmp4*-induction for 10 min. Embryos show a spectrum of phenotypes. Frequently, two optic cups are seen that are fused in the middle, synophthalmia (B, C). However, also cases of cycopia with severe pericardial effusion (*pe*) can be seen (B’ and C’). D and D’: schemes of phenotypes.The scheme in D matches with image C and D’ matches with C’. B: lateral view, head left, C, B’, D and D’: ventral view, head left. E-G: *bmp4*-induction for 7,5 min. The embryos develop ocular hypotelorism. G: scheme of phenotype, E: lateral view, head left, F: ventral view, head left, H-J: control embryo. J: scheme of phenotype, I: lateral view, head left, H: dorsal view, head left. Scale bars indicate 250 μm. The dotted lines indicate the eye domain, *y*: yolk sac. The green colour in the schematic drawings indicates eye tissue of neuroectodermal origin.

### *Cryaa* expression in *bmp4*-induced embryos

An interaction between pre-lens ectoderm and optic vesicle is fundamental for the transformation of an optic vesicle into an optic cup (Hyer et al., 2003). We next performed *bmp4* induction at 8.5 hpf for 15 minutes, 10 minutes and 7.5 minutes (**Figure 2A, scheme of experimental procedure**) and addressed the expression of *cryaa* by whole mount in situ hybridization (WMISH). *Cryaa* is a gene coding for a crystallin protein, important for lens development and function. *Cryaa* expression was regularly found in lenses of control embryos (**Figure 2B, C)**. *Cryaa* expression could also be detected in *bmp4*-induced embryos (**Figure 2D-K, metadata can be found in the supplemental table**). After induction of *bmp4* for 15 minutes, *cryaa* expression was found in patches in an anterior domain (Figure 2D and E). This domain corresponds well to the region where a lens would have been localized in a cyclopic embryo, yet no lens was formed and importantly the crypt-oculoid was not transformed into an optic cup. After induction for 10 minutes, *cryaa* expression was found expressed in lenses, if these were formed, but also in varius ectopic regions, e.g. in anterior regions of the head domain (Figure 2G and H). After induction for 7.5 minutes *cryaa* expression was found in lenses (Figure 2I and K).

**Figure 2.**
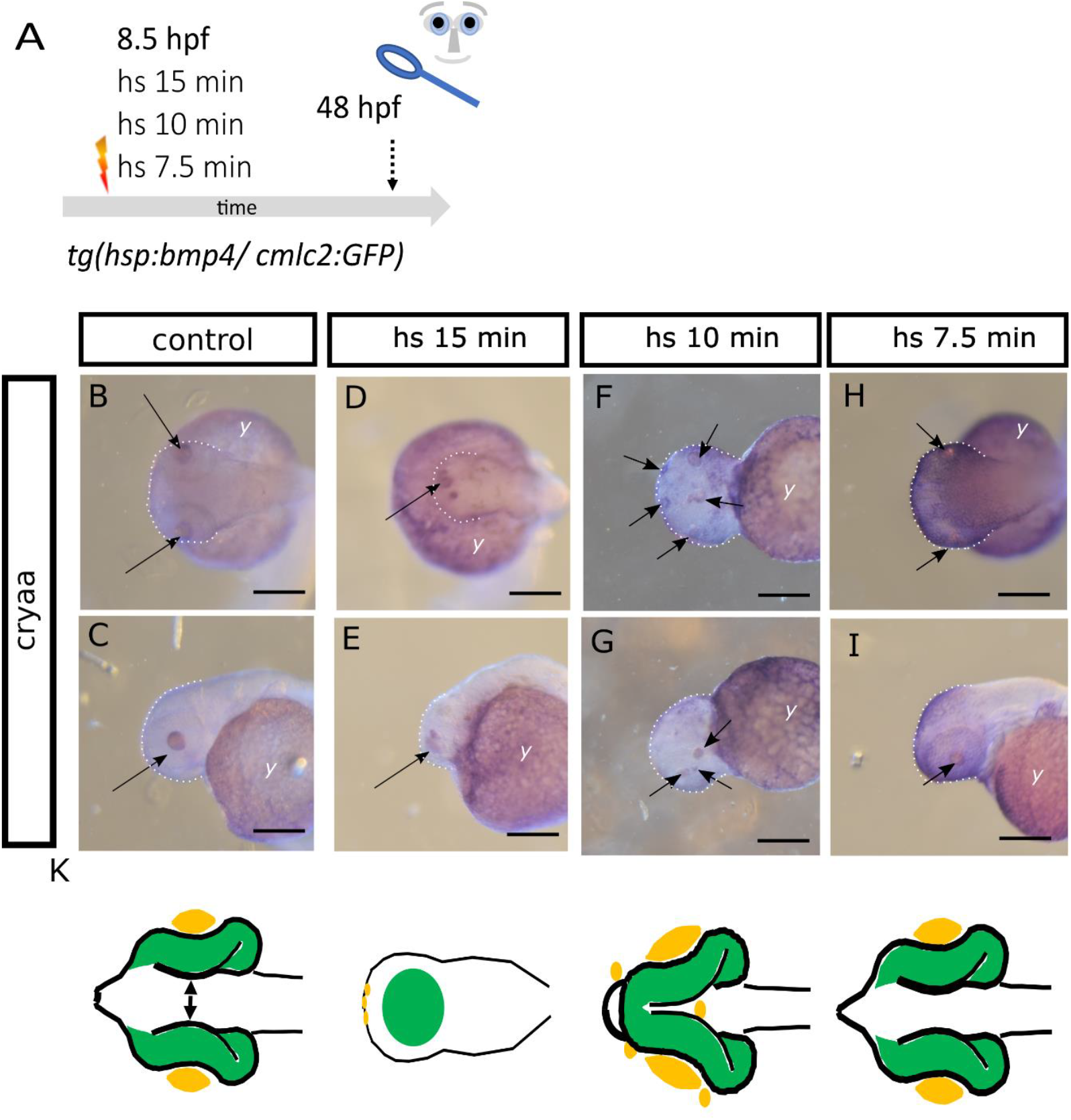
*Cryaa* expression in embryos after attenuated *bmp4* induction. A: summary of experimental procedure: embryos were heat-shocked at 8,5 hpf for 15 min, 10 minutes and 7.5 minutes and analyzed at 48 hpf. B-E: WMISH for *cryaa* at 48 hpf, B, C: control embryo, B: dorsal view, head (dotted line) left (arrow point at lenses) C: lateral view, head left (arrow points at lens). D, E: *bmp4*-induced embryo (15minutes). D: dorsal view, head left, E: lateral view, head (dotted line) left. No eyes or lenses were formed. Multiple small *cryaa*-positive domains are visible at the tip of the head (arrows in D and E). F, G: *bmp4*-induced embryo (10 minutes). F: ventral view, head left, G: lateral view, head left (arrows point at sites of *cryaa* expression). H, I: *bmp4*-induced embryo (7.5 minutes). H: ventral view, head left, I: lateral view, head left (arrows point at sites of *cryaa* expression). Scale bars indicate 200 μm. The dotted lines indicate the outlines of the heads. *y*: yolk sac. *K*: schematic representations of phenotypes. Green: eye tissue (of neuroectodermal origin), yellow: lens tissue (of ectodermal origin)

### The basal constriction of retinal progenitors is affected by *bmp4*-induction

We next aimed to further address, why the crypt-oculoid does not form an optic cup in *bmp4*-induced embryos (**Figure 2A, D, E**). During normal eye morphogenesis, individual retinal progenitors undergo a change of shape turning them from a columnar shape into a wedge shape. This process is driven by basal constriction (Martinez-Morales et al., 2009). We used the transgenic line *Tg(Ola*.*Rx2:EGFP-CAAX*,*myl7:EGFP)* to visualize the retinal progenitors with confocal microscopy. We crossed this line to the *bmp4* inducible line *Tg(hsp70l:bmp4*,*myl7:EGFP)*. To be able to address whether the change of shape of individual retinal progenitors is potentially affected by *bmp4-*induction, we delayed the *bmp4-* induction slightly. We induced *bmp4* via heat-shock at 10.5 hpf for 15 minutes (**Figure 3A, scheme of experimental design**). At this stage, eye-field splitting and optic vesicle out-pocketing has already started. Thus, it was possible to observe the effect of *bmp4* on optic cup formation independently. At 24 hpf, regular optic cups could be observed in control embryos (**Figure 3B, B’, 33 embryos**). The basal surface, directed towards the lens, showed a typical bending (**Figure 3B’**). Optic vesicle like structures were found after the delayed *bmp4*-induction (**Figure 3C, C’, 29 embryos**). The surface of these vesicles, however, remained flat (**Figure 3C’**) indicating that the shape change of the retinal progenitors was hampered. Furthermore, the vesicle like structures were not separated, but fused in the midline (**Figure 3C**). At 74 hpf the phenotype morphologically resembled the opo mutant phenotype in which the hampered basal constriction was first described (Martinez-Morales et al., 2009) (**Figure 3F,G**, **67 embryos, please see D, E as control, 39 embryos**). Besides, 5 embryos with severe degeneration were found (4 dead at 4dpf, unclear genotype) one with late onset of cmlc2: expression. We next addressed the expression of *cryaa* in the delayed induction paradigm. Unexpectedly, we did not detect *cryaa* expression after *bmp4* was induced at 10.5 hpf for 15 minutes (**Figure 3J, K, 3 embryos, please see H, I as control**). It seems possible, that the absence of lens tissue and presumably the absence of pre-lens ectoderm negatively affected optic cup formation in the delayed induction paradigm.

**Figure 3.**
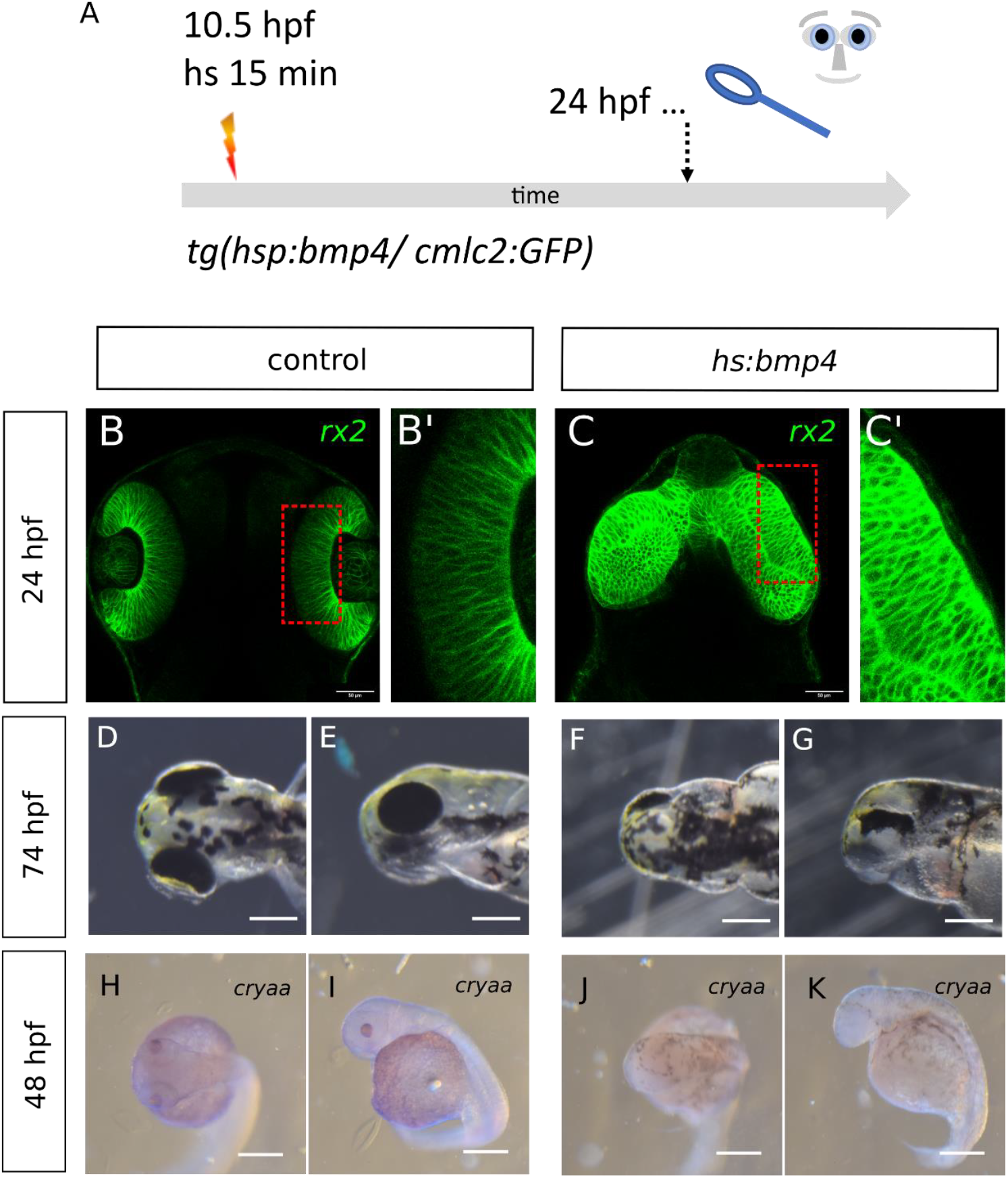
Delayed *bmp4* induction results in “flat-eyes”. A: summary of experimental procedure: embryos were heat-shocked delayed at 10,5 hpf for 15 min and analyzed 24 hpf or at later stages. B - C’: Confocal images (dorsal view) with green indicating *rx2*-positive cells at 24 hpf. C, C’: control embryo. B, B’: *bmp4*-induced embryo. Please note that the delayed induction resulted in out-pocketing of optic vesicles, which, however, were not transformed into optic cups. Scale bars indicate 50 μm. D-G: gross analysis of embryos at 74 hpf, D, E: control embryo. D: dorsal view, head left, E: lateral view, head left, F, G: *bmp4*-indced embryo. F: dorsal view, head left, G: lateral view, head left. Please note that the induced embryos show a phenotype reminiscent of a flat-eye “ojoplano” phenotype ^18^. Scalebars indicate 250 μm. H-K: WMISH for cryaa at 48, H, I: control embryo. H: dorsal view, head left, I: lateral view, head left, J, K: *bmp4*-induced embryo J: dorsal view, head left, K: lateral view, head left. Please note that the embryo shows no eyes and no visible *cryaa*-positive tissue. Scale bars indicate 250 μm.

### *Ofcc1, vsx2 and lhx2*b expressed in crypt-oculoids

The shape change of the progenitors during optic cup morphogenesis is controlled by *opo/ofcc1* (Bogdanović et al., 2012; Martinez-Morales et al., 2009). V*sx2* was found as an upstream regulator of *ofcc1* (Gago-Rodrigues et al., 2015). For the next steps of our analyses, we went back to the original, early treatment paradigm of *bmp4* induction at 8.5 hpf for 15 minutes (**Figure 4A, scheme, all embryos expressing *cmlc2:GFP* showed anophthalmia phenotype**). We addressed whether the expression of *ofcc1* and *vsx2* are affected by induction of *bmp4*. To facilitate the interpretation of expression pattern changes, a scheme of the morphological changes of the embryo was added (**Figure 4O, please see N as control**). In control embryos, *ofcc1* can be found expressed in the optic cups among many other domains (**Figure 4B, C, 5 control embryos**). *Ofcc1* expression was found in the region of the crypt-oculoid after induction of *bmp4* (**Figure 4D, E, 4 induced embryos**). In control embryos, *vsx2* expression was detected in the optic cups more specifically (**Figure 4F, G, 10 control embryos**). In *bmp4*-induced embryos, vsx2 expression was found in the head region and the crypt-oculoid (**Figure 4H, I, 10 induced embryos**). The transcription factor *lhx2b* was also found essential for the transformation of the optic vesicle into the optic cup (Porter et al., 1997). We thus also addressed *lhx2b* expression. In control embryos, *lhx2b* expression could be detected in the telencephalon, diencephalon and ventral optic cup (**Figure 4J, K, 5 control embryos**). After induction of *bmp4*, the expression of *lhx2b* could be detected in the head and the crypt-oculoid (**Figure 4L, M, 5 induced embryos**). Together, we could not detect a clear suppression of expression of neither *ofcc1, vsx2* or *lhx2b*. These findings suggest that the process of basal constriction of the retinal progenitors is more likely affected by other means than by the level of expression of these genes.

**Figure 4.**
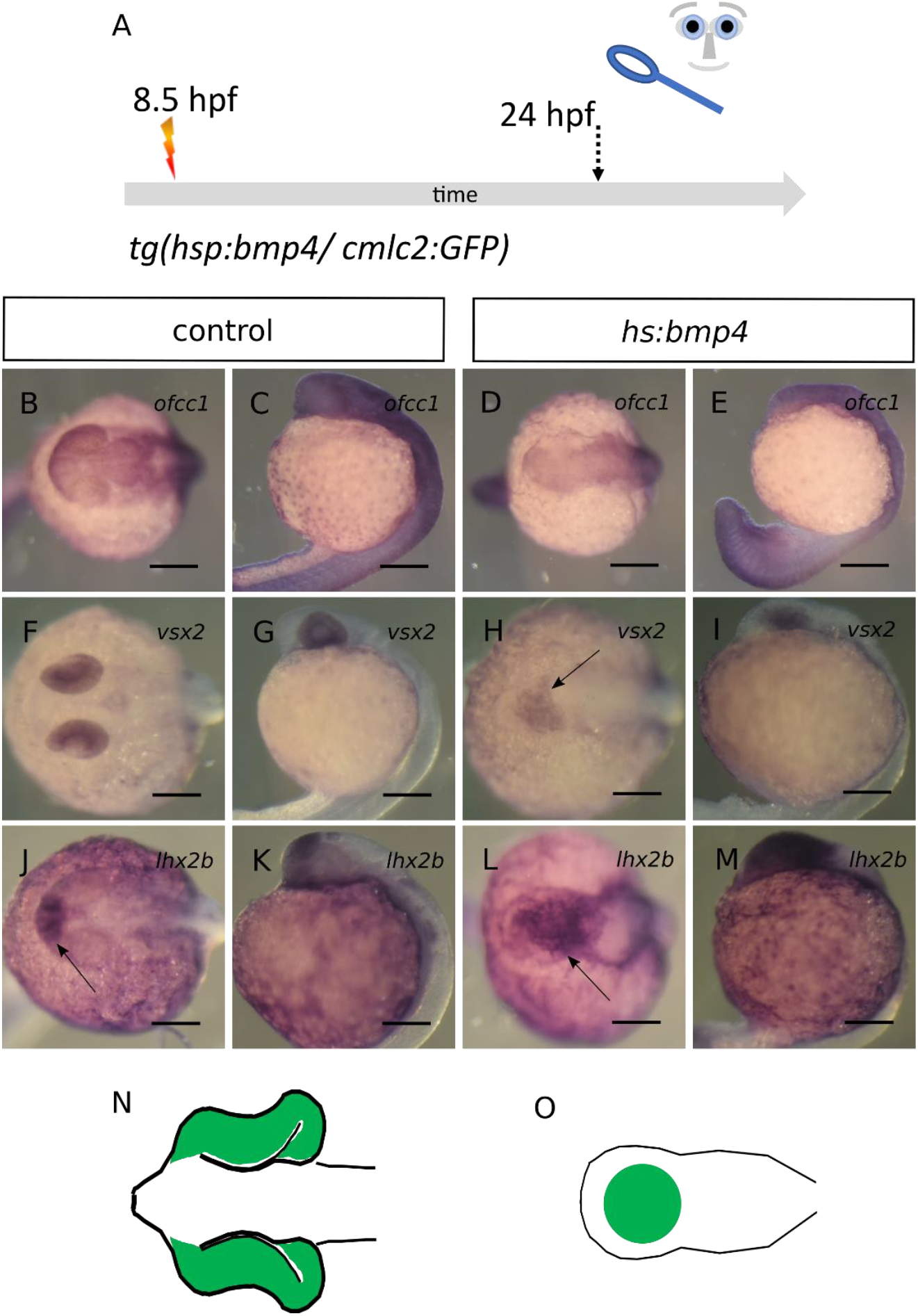
*Ofcc1, vsx2 and lhx2b are expressed in crypt-oculoids*, A: summary of experimental procedure: embryos were heat-shocked at 8,5 hpf for 15 min and analyzed 24 hpf. B-E: WMISH for *ofcc1*, B, C: control embryos, B: dorsal view, head left, C: lateral view, head left, *ofcc1* expression in optic cups and other domains, D, E: *bmp4*-induced embryos D: dorsal view, head left, E: lateral view, head left, *ofcc1* expression in the head and crypt-oculoid. F-I: WMISH for *vsx2*, F, G: control embryos, F: dorsal view, head left, G: lateral view, head left, *vsx2* expression in optic cups, H, I: *bmp4*-induced embryos D: dorsal view, head left, E: lateral view, head left, *vsx2* expression in the head and crypt-oculoid. J-M: WMISH for *lhx2b*, J, K: control embryos, J: dorsal view, head left, K: lateral view, head left, *lhx2b* expression in telencephalon, diencephalon and optic cups, L, M: *bmp4*-induced embryos L: dorsal view, head left, M: lateral view, head left, *lhx2b* expression in the head and crypt-oculoid. N, O: scheme of phenotype of control (N) and *bmp4*-induced embryo (O) “eye tissue” is marked in green. Scale bars indicate 200 μm

**Figure 5.**
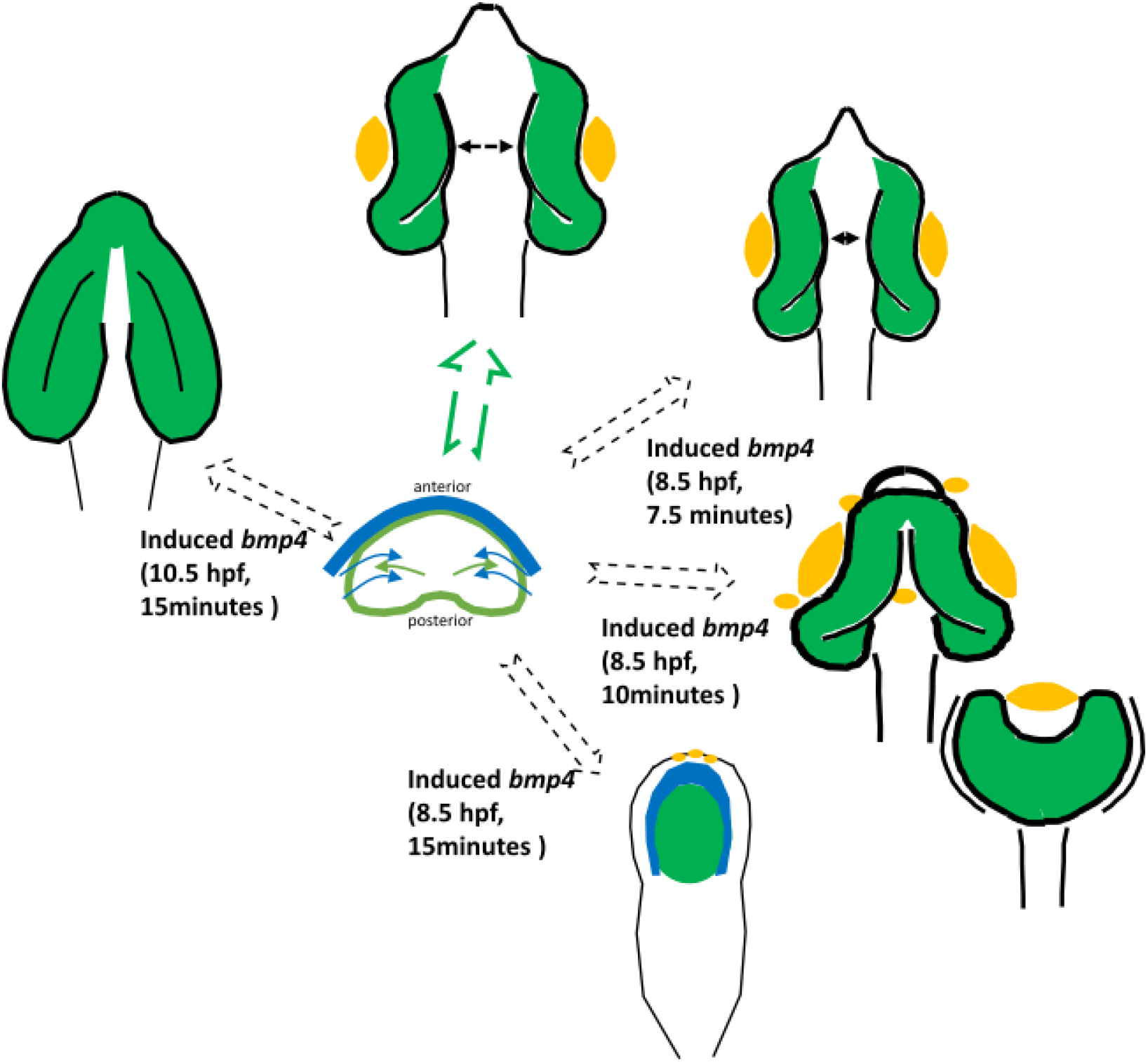
: schematic summary of findings. The eye-field (green) within the ANP (center) is split and develops (green arrow) into bilateral optic cups (top) including lenses (yellow) (arrow marks distance between optic cups). *Bmp4*-induction at 8.5 hpf for different duration results in graded phenotypes (7.5 minutes top right: ocular hypotelorism, 10 minutes middle right: synophthalmia/ cyclopia, 15 minutes bottom right: crypt-oculoid), lens tissue is marked in yellow. Delayed induction of *bmp4* (top left) hampers optic cup bending and results in flat eyes. Blue: forebrain.

## Summary and Discussion

HPE is the most frequent developmental forebrain pathology in humans (Malta et al., 2023; Matsunaga and Shiota, 1977; Pineda-Alvarez et al., 2011). The incidence of HPE is approximately 1/10000 in humans at birth. The incidence of HPE per conception is 0.4 %, determined in abortions (Matsunaga and Shiota, 1977). HPE has been graded in groups of severe forms, alobar HPE, intermediate forms, semilobar HPE and less severe forms, lobar HPE (Gomez et al., 2024). Accordingly, the classical eye phenotypes which accompany HPE can also be found as a spectrum which includes coloboma, ocular hypotelorism, synophthalmia and cyclopia (Pineda-Alvarez et al., 2011). Even though, cyclopia is often perceived to be the most extreme eye phenotype associated with HPE, also anophthalmia has been found in HPE patients (Fallet-Bianco, 2018).

Notably, the finding “anophthalmia” has to be parsed. The term anophthalmia is used in the context of the MAC spectrum (microphthalmia, anophthalmia and coloboma) (Ohuchi et al., 2019). In this context anophthalmia results from pathologies during the further development of the optic vesicles similar to the evolutionary loss of eyes in cavefish (Ohuchi et al., 2019). Here, the eye field was already split. As beforementioned, anophthalmia, however, can also be part of HPE (Fallet-Bianco, 2018). Anophthalmia in this context means that the eye field does not split and that subsequently not two optic vesicles develop. The form of HPE which underlies this eye phenotype is considered very severe. The prosencephalon is largely missing, termed pseudo-aprosencephaly (Fallet-Bianco, 2018; Schmitt and Born, 1988). The eye phenotype, anophthalmia, is thus described “most severe” in the context of HPE overall (Fallet-Bianco, 2018). Interestingly, a double Knock out of two BMP antagonists in mouse (chordin and noggin) variably affected forebrain development and cases of both cyclopia and aprosencephaly were described (Bachiller et al., 2000).

We have recently provided a new inducible HPE model in zebrafish. This is based on an induced expression of *bmp4* and a subsequent oversaturation of endogenous BMP antagonists (e.g. *nog2, chrd, fsta* and *grem2b*)(Bulk et al., 2023). The embryos were induced at 8.5 hpf, a developmental stage during gastrulation (Kimmel et al., 1995). This hampered the division of both the telencephalic domain and the eye field and resulted in anophthalmia. Notably, retinal precursors were found inside the forebrain, termed crypt-oculoid (Bulk et al., 2023).

In this analysis we aimed to investigate whether the anophthalmia phenotype could potentially be turned into less severe ocular phenotypes, e.g. cyclopia. We started from the beforementioned hypothesis that anophthalmia is more severe than cyclopia. We made use of the induction paradigm of *bmp4*. This paradigm is based on a heat-shock induced expression of *bmp4* and the phenotype anophthalmia with crypt-oculoid was observed after an induction at 8.5 hpf for 15 minutes. The level of *bmp4* expression was high enough to overcome and counteract the BMP antagonists which are expressed in the area of the ANP (Bulk et al., 2023). An inducible expression system like the heat shock promoter driven expression we used does not facilitate the control of exact expression levels. In addition, the growth factor *bmp4* can have e.g. dosage-, level- and context-depending functions. Our aim was to basically reduce the level of expression. To this end we attenuated the heat shock length. We used 10 minutes and 7.5 minutes instead of 15 minutes in order to get to an attenuated *bmp4* induction. We obtained graded phenotypes. After 10 minutes induction of *bmp4* we found cases of cyclopia and synophthalmia and after an induction duration of 7.5 minutes we found ocular hypotelorism. These findings fit well to the graded eye phenotypes which accompany HPE (Fallet-Bianco, 2018; Pineda-Alvarez et al., 2011) and to the developmental process of eye field splitting and subsequent bilateral out-pocketing of optic vesicles. These findings indicate that the experimental attenuation of *bmp4* induction worked fine and that the prominent ocular phenotypes, which can accompany HPE, can be induced in the current HPE model system. This further suggests that at least for this specific HPE model the most severe form was anophthalmia and less severe forms were cyclopia, synophthalmia and ocular hypotelorism.

It will be interesting to investigate in future projects which exact levels of *bmp4* expression result in which phenotype. Accordingly, it will be informative to dissect different signaling cascades (canonical and non-canonical) which are potentially involved and to investigate the molecular changes which result from the various *bmp* expression levels.

In our study, we next addressed why no eye was formed in this specific HPE model. We found a central anterior expression domain of the lens marker *cryaa* in the region where a cyclopic eye would have been formed, in embryos that underwent an induction of *bmp4* at 8.5 hpf for 15 minutes. This might be interpreted as part of an anlage of a cyclopic eye. Nevertheless, such a cyclopic eye was not formed in consecutive stages of development in these embryos. Bearing in mind that retinal progenitors were found inside the dysmorphic forebrain, we wanted to investigate further, why they did not form an optic cup.

We next aimed to test whether the shapes of the retinal progenitor cells are able to change after induction of *bmp4*. This change of shape during normal development is an important motor for optic cup formation, driven by *ojoplano* (*opo*) (Martinez-Morales et al., 2009). In order to be able to address this issue, we needed to slightly delay the induction of *bmp4*. This gave the eye-field more time to start to out-pocket. This way, we were able to show that the delayed induction impeded the shape changes. The phenotype which resulted resembled the *opo* mutant (Martinez-Morales et al., 2009). To our surprise, however, we were not able to find *cryaa* expressing tissue in these embryos. This indicates that the mechanism by which eye development was hampered in the delayed induction paradigm could be different. It remains elusive by what exact mechanism the optic cup formation is prevented in the early *bmp4* induction paradigm (8.5 hpf, 15 minutes). The analysis of the expression of *opo* and its upstream regulator *vsx2* was not giving further insights, neither was the analysis of *lhx2b* expression. These factors were found expressed in the dysmorphic forebrain regions of the anophthalmic embryos. This at least suggests that these factors are not functionally involved by transcriptional regulation.

Even though we cannot be sure if the impeded shape changes of retinal progenitors are functionally involved in the decision whether or not an eye is formed in the anophthalmia phenotype of the early induction paradigm, we showed that shape changes are functionally connected to BMP antagonism, overall. Our data indicate that BMP antagonism is important to allow the changes of shape of retinal progenitors. This is fitting well to the data which previously connected the “rim involution” and function of *opo* during optic cup formation (Sidhaye and Norden, 2017) and our data showing that BMP antagonism is important for optic vesicle to optic cup formation, driven by a “bilateral neuroretinal flow” (Heermann et al., 2015).

## Materials and Methods

### Zebrafish care

Adult zebrafish were kept in accordance with the local animal welfare law and with the permit from the “Regierungspräsidium Freiburg”: 35-9185.64/1.1. Fish were housed in a recirculating system at 28°C in a 12h light/ 12h dark cycle. The following transgenic lines were used: tg(hsp70l:bmp4, myl7:eGFP)(Knickmeyer et al., 2018) and tg(Ola.Rx2:EGFP-CAAX,myl7:EGFP)(Heermann et al., 2015). Zebrafish embryos were grown in petri dishes containing zebrafish medium, 0.3g/l sea salt in deionized water. Melanin-based pigmentation was inhibited, if needed for downstream applications. To this end embryos were incubated in 0.2 mM phenylthiourea.

### Heat shock procedures

Application of a heat-shock was used for heat-shock inducible transgenes. Embryos at different developmental stages were incubated in 1.5 ml reaction tubes at 37°C (heating block, Eppendorf Thermomixer). Onset and duration of the heat shock is indicated in the respective experimental setup.

### In situ hybridization

Whole-mount ISHs (WMISHs) were performed as described previously (Bulk et al., 2023; Quiring et al., 2004).

These probes were used: *vsx2* (Eckert et al., 2020), *ofcc1* F: GATGCTGCCAAGCTCTACTGG; R: TTCATCCTCTCGTGTTGCTCAT, *lhx2b* F: ATGCCTTCAATCAGCGGG, R: TCAGAAGAGGCTGGTTAAGG, probes for *cryaa* were designed cloning free (Hua et al., 2018). Sequences are: *cryaa* F: CCAACACCCTTGGTTCAGAC, *cryaa* R: GTAACAGGGATGGTGCGATCT

### Image processing

Images were recorded via microscopy (Nikon, binocular) and images were subsequently edited for presentation using the software ImageJ (Fiji) (Schindelin et al., 2012) and Inkscape (Inkscape Project. (2020). *Inkscape*. Retrieved from https://inkscape.org).

## Acknowledgments

We want to thank all members of the Heermann lab for insightful discussions. We want to express special gratitude to Ute Baur for great technical assistance. Furthermore, we want to thank Eleni Roussa and Klaus Unsicker for support of the Heermann lab. For support in regards to publication we want to thank the Open Access Publication Fund of the University of Freiburg.

## Figures and Figure Legends

**Figure 1 supplement.**
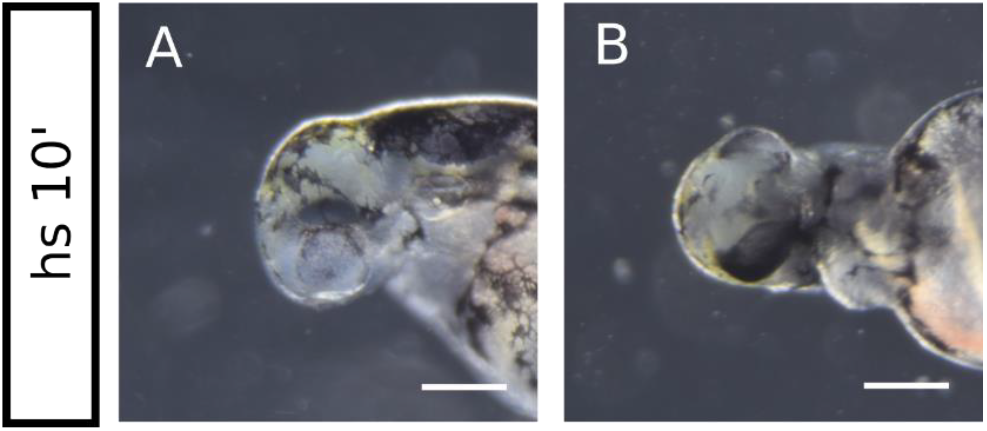
Spectrum of ocular phenotypes. A and B: *bmp4*-induction for 10 min. Optic cups are fused in the middle resembling a strong case of synophthalmia. A: lateral view, head left, B: ventral view, head left. Scale bars indicate 250 μm.

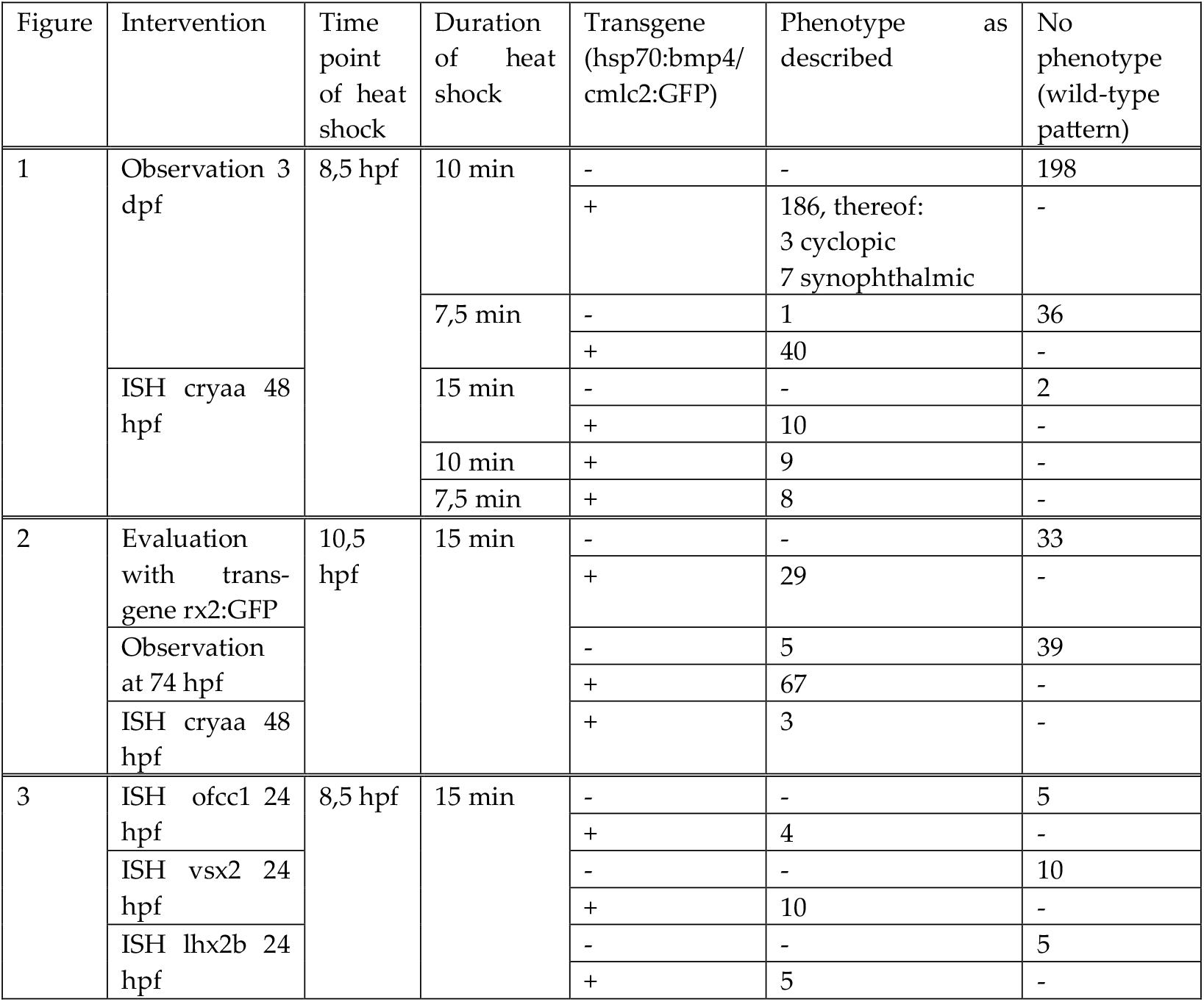

